# A conceptual framework for revealing minor bacterial signals in microbiome data through guided data transformation

**DOI:** 10.1101/2025.05.31.656121

**Authors:** David Martin, Pierre Houedry, Frederic Derbré, Valerie Monbet

**Affiliations:** IRMAR - UMR CNRS 6625, University of Rennes, Rennes, France; Laboratory “Movement Sport and Health Sciences”, University of Rennes 2/ENS Rennes, Rennes, France

## Abstract

The microbiome is a rich source of biological data that offers promising insights into personalized medicine. However, inferring host health from gut bacterial composition using statistical analytical methods remains a challenge. Here, we show that groups of bacterial species with high abundance and variance (referred to as dominant bacterial signals and often associated with enterotype) exert a disproportionately large influence on microbiome analyses, hiding the contribution of less expressed species (referred to as minor bacterial signals). To address this limitation, we propose a guided data transformation highlighting minor bacterial signals while minimizing the impact of dominant bacterial signals on microbiome statistical analyses. This transformation (i) leads to alternative clustering more closely associated with host health and (ii) helps to improve the performance of supervised machine learning algorithms in high-dimensional settings (*n* ≪ *p*). Applying to a real dataset, our results suggest that dominant bacterial signals may act as a confounding variable to predict host health.

## 1 INTRODUCTION

The gut microbiota is increasingly recognized as a rich source of biological data that offers critical insights into host health, including information on pathological conditions such as cancer, diabetes, and other metabolic disorders (*1*). In this context, numerous European and American companies have entered the lucrative personalized microbiome analysis market, driven by both preventive and therapeutic ambitions. Although many studies have developed statistical analyses of microbial composition data derived from metagenomics sequencing (*2*), substantial research is still needed before scientifically validated health claims can be established based on the gut bacterial ecosystem (*3*).

A widespread approach that involves identifying patterns, structures, or latent features within the data is known as unsupervised analysis. These methods do not rely on predefined labels. In microbiome research, unsupervised approaches typically utilize clustering or dimensionality reduction techniques, which create a simplified and synthetic representation of gut bacterial composition (*4*).

In this context, the enterotype classification system has emerged as the predominant clustering framework in gut microbiota analysis and is still widely applied (*5, 6, 7*). The original enterotype model categorizes individuals into three distinct groups, each defined by the dominance of a specific bacterial genus: Prevotella, Bacteroides, or Ruminococcus. Enterotypes are conceptually defined as ffdensely populated regions within a multidimensional space of community compositionff (*8*). The enterotype-based clustering framework reduces the complexity of gut microbiota data by partitioning it into a small number of interpretable clusters. Consequently, this stratification is now considered a potential guide for personalized interventions targeting host health (*9*). Since its introduction, additional enterotypes have been proposed, often varying depending on the population or species studied (*10*). However, this approach is also criticized for lacking well-defined cluster boundaries, raising concerns about the robustness of enterotype classifications (*11, 12*).

Moreover, this method, characterizing differences between microbiomes, is driven by species with the highest variance. However, it has been shown that species variance is positively correlated with their abundance, which introduces bias into ecological and microbiome analyses (*13, 14, 15*). Here, the species showing a high abundance and variance are referred to as the dominant bacterial signals, while species with low abundance and variance are referred to as the minor bacterial signals.

On the other hand, analytical approaches used to predict health status based on the gut microbiome are typically classified as supervised. Supervised analysis refers to methods that rely on labeled datasets, in which the outcome variable, such as the host health variable, is known (*16*). These approaches aim to learn a mapping between the input characteristics and the target variable, often using machine learning methods combined with cross-validation techniques to optimize predictive performance (*17*). However, the known relationship between the abundance and variance of species, where highly abundant species exhibit a high variance, can also bias supervised analyses (*14*) and also the interpretability of the model (e.g., species or strain detection). The implications of this relationship for both characterizing the gut microbiome and predicting host health appear underexplored.

In this study, we aim to address this gap by focusing on the challenge of inferring host health from gut bacterial composition, using both unsupervised and supervised analytical methods. Using public datasets from healthy individuals and patients with irritable bowel disease (*18*), we demonstrate that groups of species characterized by high abundance and variance (referred to as dominant bacterial signals) primarily drive clustering patterns and are preferentially selected as important features in supervised classification. Conversely, species with lower abundance and variance (referred to as minor bacterial signals) are overshadowed by the dominant bacterial signals, despite exhibiting relevance in characterizing host health.

Then, we address this observed bias by introducing a guided data transformation. It aims to preserve minor unknown signals and decrease the impact of known dominant bacterial signals (e.g., enterotype) on subsequent analyses. By developing an original simulation pipeline, we show that the introduced transformation helps to reduce the strong effect of the dominant bacterial signals by improving the performance of machine learning algorithms in high-dimensional settings (*n* ≪ *p*). In that case, we also display that the interpretability of the models is improved. Finally, the guided transformation has been applied to other public datasets, highlighting that enterotype could act as a confounding variable to predict host health.

## 2 RESULTS AND DISCUSSION

### 2.1 Reference dataset used for demonstration and numerical experiments

The reference dataset, used for the following demonstration, is registered in Bio-project n°PRJEB1220 (*18*). The dataset consists of 396 samples, each characterized by the expression levels of 606 microbial species and an associated binary host health label: 0 for healthy samples (HLT) and 1 for samples diagnosed with Irritable Bowel Disease (IBD).

### 2.2 The dominant bacterial signals drive the clustering of gut microbiome

In gut microbiome studies, pairwise (dis)similarity is frequently assessed using the Bray-Curtis metric. The clustering based on this (dis)similarity matrix reveals three distinct clusters (Figure 1. A.1-2), closely resembling the enterotype-based clustering framework. Specifically, the yellow cluster includes microbiomes with a high abundance of species from the *Prevotella* genus, while the light and dark pink clusters are primarily composed of microbiomes dominated by species from the *Bacteroides* genus.

**Figure 1:**
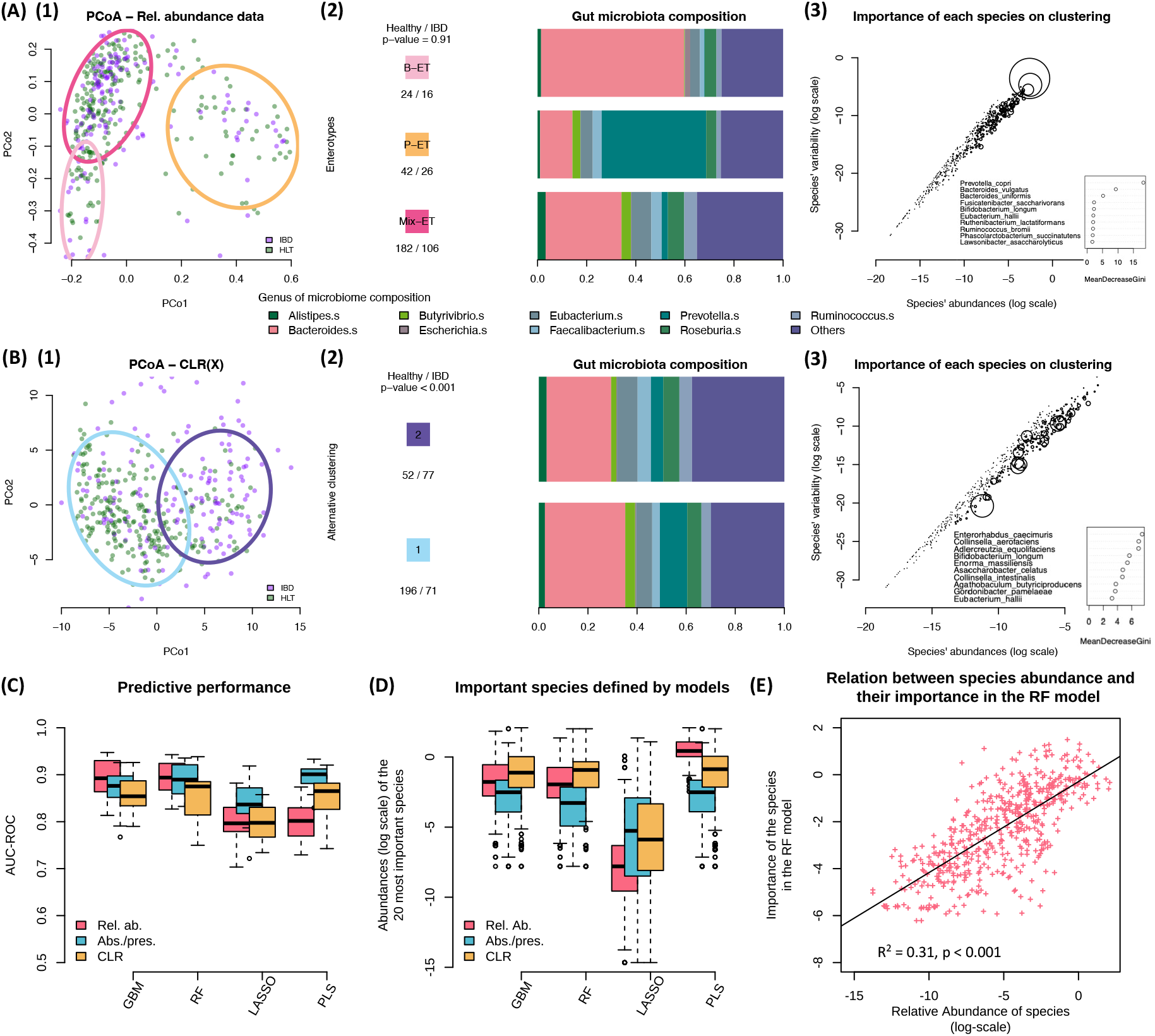
Observed problem: the dominant bacterial signals, close to the enterotype clustering system, drive the analyses of microbiome data, hiding minor bacterial signals. (**1**) Principal Coordinates Analysis and the 80% confidence ellipse of the clusters (in colored circles) are presented. (**2**) Distribution of each host health condition per cluster. The *p*-value of the *χ*^2^ between the clustering and the host health is computed. Then, a barplot of the microbiome composition, at the genus level, for each cluster is shown. (**3**) Abundance-variance relationship at the species level, illustrating the influence of each species on the clustering. Circle size reflects the species’ contribution to the clustering. The 10 most influential species (i.e., the largest circles) are highlighted along with their corresponding mean decrease in Gini index. The following demonstration has been applied to both relative abundance data (**A**) and centered log-ratio (CLR) transformed data (**B**). (**C**) The predictive performance is evaluated using the Area Under the Receiver Operating Characteristic Curve (AUC-ROC). Four distinct machine learning models are computed (Gradient Boosting Machine (GBM), Random Forest (RF), Lasso regression (LASSO), and Partial Least Squares (PLS)). (**D**) The distribution of the mean abundances (log-transformed) of the 20 most important species defined by distinct machine learning models. (**E**) The strength of the relationship between species importance defined by the RF model and species abundance is assessed using the coeffcient of determination (*R*^2^). IBD refers to Inflammatory Bowel Disease microbiomes, and HLT refers to Healthy microbiomes.

A Random Forest algorithm is applied, where the target variable corresponds to the clustering assignment, a qualitative variable, and the explanatory variables are the bacterial composition of the gut (*13, 15*). The importance scores of species derived from the model are interpreted as their respective contributions to the clustering. Subsequently, we examine the abundance–variance relationship at the species level, which confirms previous observations. To visually represent species contributions, we use circles of varying sizes, with larger circles indicating greater influence. In particular, we highlight the 10 most influential species (i.e., those with the largest circles), along with their corresponding mean decrease in Gini index (Figure 1.A.3). These results underscore a strong association between species characterized by high abundance and variance, and their impact on clustering. Notably, *Prevotella copri, Bacteroides vulgatus*, and *Bacteroides uniformis* (which are the three species with the highest abundance and variance, in this dataset) appear to drive the clustering patterns (Figure 1.A.3).

Consequently, and unsurprisingly, we show that clustering and projection analyses based on the BrayCurtis metric are primarily influenced by dominant signals (i.e., highly expressed species), whereas minor bacterial signals (i.e., less abundant species) are not fully captured.

This issue can be addressed: by modifying the (dis)similarity/distance metric between samples. For example, the UniFrac distance leads to an alternative clustering by decreasing the importance of phylogenically close species, such as *Prevotella* and *Bacteroides* (*19*); by applying transformation on data, such as log-ratio-based transformation (*20*). Log transformation methods have already been identified to be effcient by decreasing the variability of highly abundant species (*14*); by converting all positive abundances to a value of one. The presence/absence matrix is also considered relevant to decrease dataset complexity.

In this context, we demonstrate that clustering based on distinct data transformations (that is, log-transformation, UniFrac, absence/presence) results in alternative clustering exhibiting stronger association with host health. Specifically, the second alternative cluster (represented by the purple circle) is predominantly composed of IBD samples, whereas the first alternative cluster (represented by the blue circle) consists primarily of healthy samples, as it illustrated in Figures 1.B.1-2 and S1.A-B.1-2.

We confirm the association between alternative clustering and host health by computing the p-value of the *χ*^2^ test, where the null hypothesis (*H*_0_) states that host health is independent of clustering, which appears lower than 0.05. This indicates a significant relationship between alternative clustering and host health (Figures 1.B.2 and S1.A-B.2). Interestingly, we show that these alternative clusters are influenced by minor bacterial signals (which are species with pretty low abundance and variance) (Figures 1.B.3 and S1.A-B.3).

Together, our analyses demonstrate that dominant bacterial signals predominantly drive current microbiome clustering, while data transformation appears to be a relevant approach to capture minor bacterial signals, which appear relevant to highlight host health patterns. Based on these findings, we hypothesize that increasing the importance of minor bacterial signals, which are typically overlooked, may provide more accurate information on host health.

### 2.3 The dominant bacterial signals represent the most important features in predictive classification: the case of high dimensionality

In the following section, we focus on supervised analyses that are affected by the curse of dimensionality, especially because clinical studies often present limited sample sizes. However, these findings can be extended to larger datasets. Now, let **X** = (*X*_*ij*_) ∈ ℝ ^*n*^ ^×*p*^ be the matrix of the microbiome dataset, where *n* and *p* are, respectively, the number of samples and species.

A large portion of microbiome research involves supervised analysis, which refers to methods that rely on labeled datasets to predict an outcome variable based on microbiome composition. These approaches pose unique analytical challenges due to the inherent sparsity and high dimensionality of microbiome data (*16*). Such challenges are particularly pronounced in clinical studies, where limited sample sizes can significantly affect the robustness and generalizability of predictive models. This is commonly referred to as the *n* ≪ *p* scenario, where the number of features (*p*) far exceeds the number of samples (*n*), and the amount of information available often governs the predictive performance. The information contained in a dataset, indicated as *I* (**X**), can be defined by the total variance of microbial species:

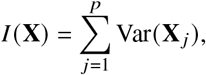

where **X** _*j*_ denotes a variable (i.e. a species).

In general, the greater the amount of information in a dataset, the better a model can fit the data. However, in *n* ≪ *p* cases, heterogeneity in intestinal microbiota composition between samples can hinder accurate prediction of host health due to a lack of consistent bacterial signatures.

To mitigate this issue, applying a logarithmic transformation or converting the dataset into a presence/absence matrix may theoretically enhance predictive performance by reducing overall information content and minimizing the influence of dominant, highly variable characteristics (*14, 21*).

On the other hand, extensive research has shown that decision tree-based models are particularly well suited for metagenomic data, largely due to 1) their non-parametric nature, 2) their ability to catch local patterns, and 3) their robustness in handling high-dimensional datasets (*22, 23, 24*). In addition, the interpretability of the model is essential in biological sciences.

For this demonstration, we focus on the cases where *n* ≪ *p*. The training set consists of 80 randomly selected samples from the reference dataset. Predictive performance is evaluated on a separate set of 100 additional randomly selected samples from this dataset. The target variable is the host health, either IBD or HLT. We train four different machine learning models: Gradient Boosting Machine (GBM) and Random Forest (RF), Lasso regression (LASSO) and Partial Least Squares (PLS).

Surprisingly, our results, demonstrate that models fitted to log-transformed data reduce predictive performance (Figure 1.C). Our findings raise a key limitation of the log transformation lying in its unguided nature. Indeed, the log-transformation is applied uniformly across all species and may reduce the variability of all abundant species linked to host health, potentially resulting in the loss of valuable biological information. Moreover, models trained from log-transformed data seem to rely more on species with higher abundance compared to models trained in relative abundance data (Figure 1.D). Then, the predictive performances of models trained from the absence/presence matrix do not outperform other transformation methods. However, these models base their predictions on less-expressed species. Notably, the performance of the PLS model fitted on the absence/presence matrix is as effective as the performance of the Random Forest model fitted on relative abundance data.

Our result also supports the relationship between species importance derived from the RF fitted on the relative abundance data and species abundance confirms that species importance in the machine learning model is dependent on species abundance (Figure 1.E).

Consequently, we show that standard machine learning models mainly involve on dominant bacterial signals and generally do not take into account minor bacterial species, which could be more relevant in some cases. We hypothesize that the presence of minor bacterial signals may lead to a more accurate insight into the host health, whatever the type of analysis.

### 2.4 Modeling the observed problem and proposed postulates

The problem under investigation is modeled by assuming that the microbiome data are structured into *M* ≥ 2 groups of species (dominant/minor bacterial signals). The groups of species are not necessarily disjoint and are unknown.

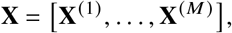

where **X**^(*m*)^ is composed of several species 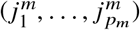 and can be characterized by its information:

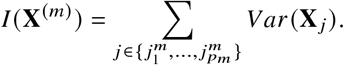

We postulate that there exist dominant bacterial signals and minor bacterial signals. It can be translated as follows:

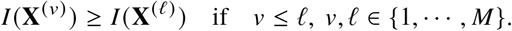

In other words, the composition of the gut microbiota includes a group of species, represented by **X**^(1)^, which exhibits higher information than another bacterial group within the same sample (e.g. **X**^(2)^, **X**^(3)^, …, **X**^(*M*)^). The goal of the proposed method is to remove a part of the information contained in **X**^(1)^. We hypothesize that this will enable, in a subsequent step, the emergence of alternative results.

### 2.5 Guided data transformation

The proposed transformation is based on two steps described in the following. The first step helps to synthesize the variability between samples related to the dominant species. It is done through a clustering. The second step helps to remove some mean information. It consists of computing residuals of linear models. The procedure is also summarized in Algorithm 1 and has been implemented in the function `estimate n residuals`in the Python module named `BiomeSampler.`

#### 1) Estimation of a clustering of the observations related to the dominant bacterial signals

we add an extra assumption. We assume that the observations are driven by a clustering such that

1. *Z* is a qualitative latent variable which takes this values in {1, ···, *K*} with *a priori* distribution *P*(*Z* = *k*) = π_*k*_, ∀*k* ∈ {1, ···, *K*}.
2. 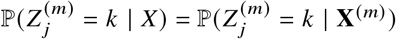.

In practice, the groups of variables **X**^(*m*)^ are unknown. However, let us remark that assumption (ii) implies that the clustering obtained from the observations **X** is the same as the clustering that would be inferred from **X**^(1)^.

The Hierarchical Ascendant Clustering (HAC) on the pairwise dissimilarity matrix (using non-Euclidean metrics such as Bray Curtis or Jaccard) of compositional data **X** leads to a clustering, i.e. estimation of 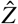. It is mainly driven by the group of species with the highest degree of information (i.e. **X**^(1)^, dominant bacterial signals). It corresponds, in most gut microbiota cases, to the clustering related to enterotypes (as illustrated in Figure 1). In practice, the optimal number of classes is determined by selecting the number that maximizes the silhouette coeffcient.

#### 2) Removing information related to the dominant bacterial signals

For each variable of {*X*_1_, ·· ·, *X*_*p*_}, the linear effect of 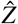 is removed as follows:

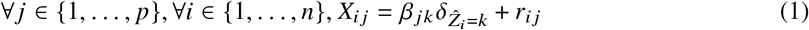

Remark that the estimation of model (1) leads to residuals *r*_*j*_ that are centered. Furthermore,

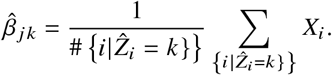

It means that 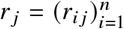 is a counter part of *X*_*j*_ where a mean effect of the dominant species is removed. We focus our next analysis on the residuals **r** = (*r*_*ij*_)_*i*_=1,···,*n, j* =1,···, *p*, which are referred to as the guided transformed data. They can be interpreted as the part of the data that is not associated with 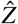. The effect of the guided transformation applied to *Bacteroides Uniformis* is illustrated in Figure S2.

Note that this step (removing information) can be applied to relative abundance data and/or to transformed data.

##### Algorithm 1

Guided transformation

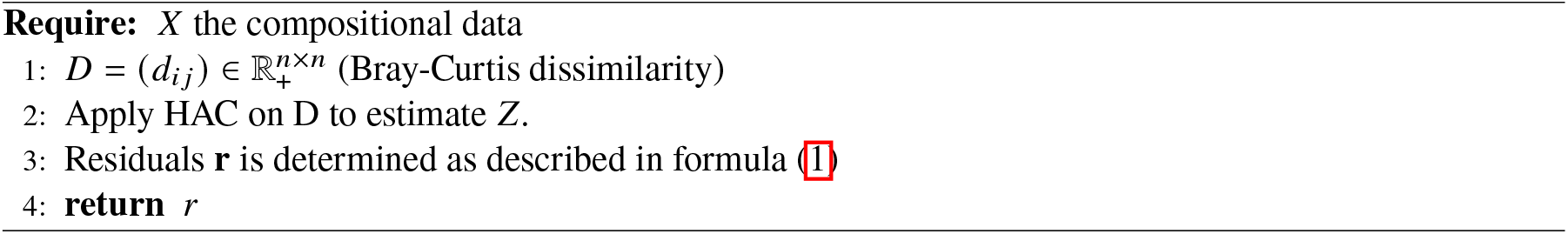

### 2.6 Overview of the numerical experiments

The next two sections test the guided transformation across various scenarios. It answers to the central hypotheses: (i) the various data transformations facilitate the formation of clusters associated with host health by reducing the influence of dominant bacterial signals, and (ii) the various data transformations improve predictive performance in a *n* ≪ *p* problem by selecting minor bacterial signals.

The reference dataset is used to simulate the data in order to validate the proposed methodology. We aim at quantifying the robustness and informativeness of microbiome-derived features under varying assumptions of signal strength and sample size. For that, the host health target variable (*Y*_sim_) is computed as a function of the minor bacterial signals, and we propose three scenarios in which the dependence between the dominant bacterial signals and the target variable *Y*_sim_ is categorized as low, medium, or high. This is managed by introducing noise into host health pattern. Those situations are tested in different dimensions where the number of samples varies (*n*_*sim*_ ∈ {75, 150, 300}). Fifty datasets have been simulated. The details of the experimental design and methods used to simulate the microbiome data are provided in the Materials and Methods section and the Figure 5.

For each condition, we assessed (i) the clustering performance by evaluating the association between transformed data clusterings and host health, and (ii) the predictive performance of machine learning models trained to predict host health status from microbiome features.

Before applying the proposed methodology, we have to check that the simulated data have a similar structure as the one observed in the reference dataset. In particular, the simulated data should present three distinct enterotypes, each driven by species predominantly from the genus *Prevotella* or *Bacteroides*. As anticipated based on our simulation design, samples assigned to the enterotype dominated by *Prevotella* species were those most frequently associated with Inflammatory Bowel Disease (IBD). It is confirmed by a statistical test where the null hypothesis (*H*_0_ :)‭the composition of the enterotype is independent of the simulated host health status‭ is rejected with a *M*-value *<* 0.001 (see Fig. 6).

### 2.7 Data transformations facilitate the formation of clusters associated with host health by focusing on minor bacterial signals

For each subset of experimental conditions and type of data, we assessed clustering performance. A confusion matrix is constructed between the host health status and the clusters. Then, *χ*^2^ test is applied to this matrix to evaluate the degree of association. The null hypothesis (*H*_0_) states independence between clustering and host health.

Here, we show that the clustering performance derived from transformed data remains effective in detecting host health, whatever the experimental scenario, especially when the influence of dominant bacterial signals on *Y*_sim_ is high (Figure 2.A). Furthermore, our results demonstrate that the clustering applied to transformed data is more influenced by minor bacterial signals than the clustering applied to data of relative abundance (Figure 2.B). Data transformations appear essential to capture minor bacterial signals in microbiome data.

**Figure 2:**
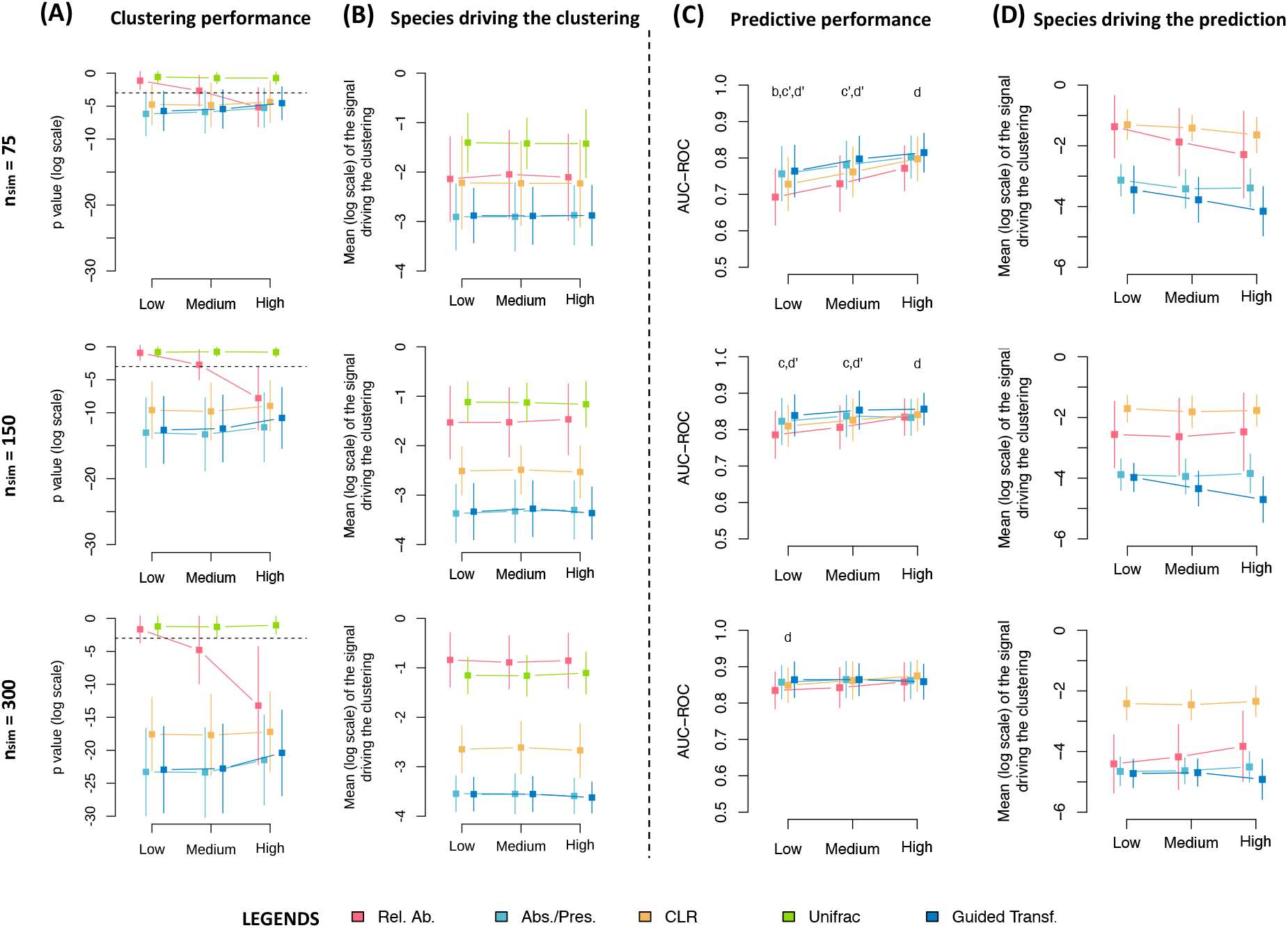
Guided transformation of microbiome data enhances clustering performance and prediction accuracy in high-dimensional, low-sample-size settings (*n* ≪ *p*). Three simulation scenarios are considered, where the influence of dominant bacterial signals on the simulated host health outcome (*Y*_sim_) is low, medium, or high. These scenarios are evaluated across different sample sizes: *n*_sim_ ∈ {75, 150, 300}. (**A**) Clustering was performed for each scenario. Performance was evaluated using a *χ*^2^ test on the confusion matrix between cluster assignments and *Y*_sim_. Distributions of log-transformed p-values across 50 simulated datasets are shown. The dashed line indicates the log 0.05 threshold, below which the clustering is statistically relevant with host health. (**B**) The bacterial species driving the clustering were identified using Random Forest models. The distribution of the mean abundances (log-transformed) of the top 20 most important species is shown across simulations. Statistical comparisons are omitted, as the patterns are visually distinct. (**C**) Predictive performance was assessed using the AUC-ROC on test sets. Letters denote statistically significant differences (*P <* 0.05) between the relative abundance data and: centered log-ratio (CLR) transformation (*k*), absence/presence transformation (*c*), and guided transformation (*d*). A prime symbol (^′^) indicates additional significant differences between CLR and other transformations. (**D**) The distributions of the top 20 most important species for predicting host health are shown. As in (B), statistical marks are omitted due to visually discernible trends. The guided transformation has been applied to the Absence/Presence matrix.

Consequently, the unsupervised analysis derived from various transformed data takes into account the presence of minor bacterial signals, leading to a more accurate insight into the simulated host health (*Y*_sim_). We validate that in a situation where the dominant signal has a strong effect on host health, the unsupervised analysis of transformed data remains effcient by focusing its analysis on minor bacterial signals. However, this guided data transformation does not lead to a higher clustering performance than other data transformations.

### 2.8 The guided data transformation improves predictive performance in an *n* ≪ *p* problem by selecting minor bacterial signals

The scenario and the methods used in this section are the same as the previous section. For each subset of experimental conditions, a Random Forest (RF) model was trained on a training set of varying sizes (*n*_train_ ∈ {75, 150, 300}) and evaluated on a fixed test set (*n* _test_ = 100). Predictive performance was assessed using the area under the receiver operating characteristic curve (AUC-ROC). Additionally, we examined the distribution of the 20 most important bacterial species contributing to the prediction of the host health variable.

Our results demonstrate that training RF models on guided transformed data and presence/absence data improves predictive accuracy in settings where the curse of dimensionality is more severe (*n*_train_ = 75, Figure 2.C). Notably, only RF models trained on guided transformed data consistently outperform those trained on relative abundance data when dominant bacterial signals exert a strong influence on host health (*n*_train_ = 75, Figure 2.C). These findings support the hypothesis that reducing the total information improves predictive performance in high-dimensional small-sample datasets. In addition, models trained on guided transformed data are primarily influenced by minor bacterial signals (Figure 2.D).

RF algorithms trained on logarithmically transformed data show predictive performance comparable to the ones trained in other transformed data sets, but do not outperform those using relative abundance data (Figure 2.C). In addition, these models are predominantly based on dominant bacterial signals, even though host health was simulated based on minor bacterial signals (Figure 2.D). Consequently, the interpretability of RF algorithms trained on log-transformed data appears limited.

Importantly, the benefit of data transformation diminishes as training sample size increases (Figure 2.C).

In larger datasets, RF algorithms trained on relative abundance data increasingly leverage low-abundance species, indicating that minor bacterial signals are essential for accurate prediction of host health.

In summary, our results show that data transformations (particularly guided transformation) improve predictive performance. These transformations achieve this by focusing the analysis on biologically relevant minor bacterial signals. Building on these findings, we now evaluate the guided transformation on real datasets. The first application addresses a high-dimensional (*n* ≪ *p*) predictive problem, while the second illustrates how guided transformation can reveal enterotypes as potential confounding factors.

### 2.9 The decrease of the total amount of information enhances predictive performance of fat oxidation in humans

In this first application of our guided data transformation, we aim to develop a predictive model to classify fat oxidation (FO) rates in humans, to determine whether individuals exceed or fall below the threshold of 0.4 g/min, using gut microbiota composition as a predictor (*25*). The 0.4 g/min threshold for FO was selected because it has previously been identified as an effective cut-off to discriminate between populations with poor versus normal metabolic flexibility (*26, 27*). A detailed description of the dataset is provided in the Materials section (*28*).

An initial clustering analysis based on relative abundance data revealed two distinct enterotypes: one associated with the genus *Bacteroides* and a second with species belonging to the *Prevotella* genus, represented respectively by pink and yellow circles in Figure 3.A. A χ^2^ test, which states that FO is independent of the clustering, yielded a p-value of approximately 0.006 (Figure 3.B). It indicates a significant relationship between dominant microbial signals (i.e., enterotype-defining species) and fat oxidation. We confirm that the most influential species in the clustering are those with the highest abundance and variance (Figure 3.C), including *Prevotella Copri* and *Bacteroides Uniformis*.

**Figure 3:**
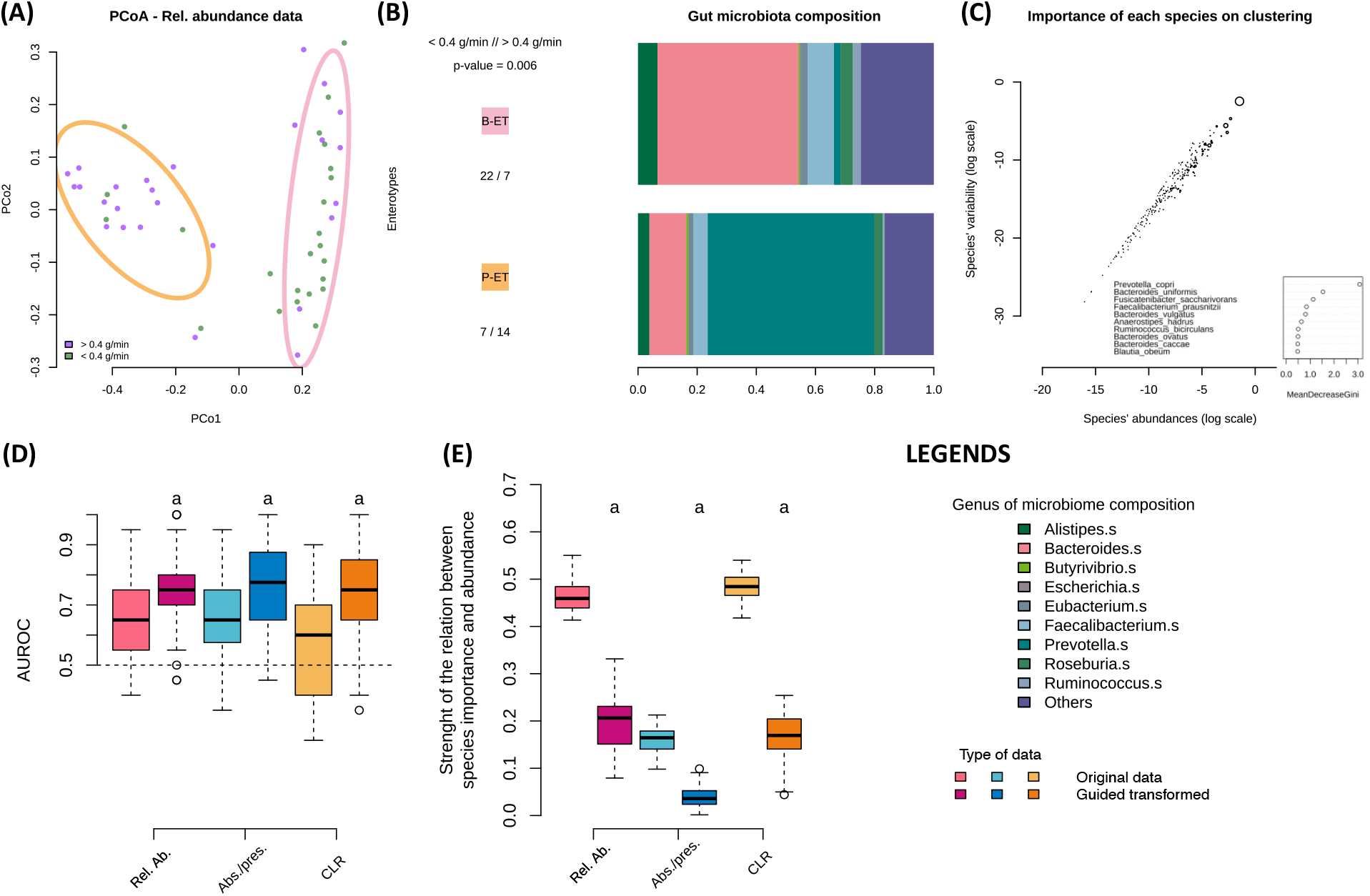
(**A**) Principal Coordinates Analysis (PCoA) with 80% confidence ellipses for each cluster (colored circles).(**B**) Distribution of host health status across clusters. The p-value of the *χ* ^2^ test assessing the association between clustering and host health is reported. A bar plot showing the microbiome composition at the genus level for each cluster is also presented. (**C**) Abundance-variance relationship at the species level, illustrating the influence of each species on the clustering. Circle size reflects the species’ contribution to the clustering. The 10 most influential species (i.e., the largest circles) are highlighted along with their corresponding mean decrease in Gini index. (**D**) Predictive performance of Random Forest models across different transformation strategies, with and without guided transformation. (**E**) Strength of the linear relationship (expressed as *R*^2^) between species abundance and feature importance in each Random Forest model. The symbol *a* indicates statistically significant differences between the original data and its guided transformation. The guided transformation has been applied to all types of data.

To investigate this further, we compared the predictive performance of Random Forest (RF) models trained on relative abundance data with those trained on various transformed datasets, including guided transformed data. The guided transformation was applied to each type of data as defined in Equation (1). RF models were trained using 80% of the samples, and predictive performance was evaluated on the remaining 20%. This procedure was repeated twenty times to estimate the distribution of AUC-ROC values.

RF models trained on both relative abundance and presence/absence data yielded mean AUC-ROC values of approximately 0.65 and 0.66, respectively. Applying the guided transformation to either of these data significantly improved the AUC-ROC by an average of 0.05 points (*p <* 0.05, Figure 3.D). These results support our hypothesis that reducing the overall information can enhance the predictive accuracy of supervised algorithms, especially in small-sample, high-dimensional contexts.

Furthermore, we assessed the interpretability of the model by computing the coeffcient of determination (*R*^2^) between species abundance and their importance in RF models. We found that models trained on guided transformed data relied more on minor bacterial signals compared to other data (Figure 3.E). These findings suggest that incorporating minor bacterial species into the models could lead to improved predictive performance.

In conclusion, these findings provide evidence that, in *n* ≪ *p* scenarios, dominant bacterial signals can obscure more informative minor bacterial signals. Guided transformation effectively mitigates this issue, leading to both improved predictive performance and interpretability in predicting human fat oxidation based on gut microbiota composition.

### 2.10 The dominant bacterial signals appear to act as a confounding factor in predictive models of Ulcerative Colitis

In this second application of our guided data transformation, we aim to develop a predictive model to classify host health, specifically whether they have ulcerative colitis (UC) or not, using gut microbiota compositions as predictors. The dataset comprises 12 public datasets. All gut microbiomes were curated in a previous study (*29*), resulting in a unified dataset of 1328 samples including 925 bacterial species.

We replicated the findings of Wu et al. (2024) (*29*), confirming the presence of three distinct enterotypes within the dataset (Figure 4.A-B). The *Bacteroides* enterotype (ET-B), depicted in red, comprises a comparable number of healthy and UC samples. In contrast, the *Clostridium* enterotype (ET-C), represented in yellow, consists almost exclusively of UC samples. The third enterotype is characterized by a diverse assemblage of genera, predominantly *Blautia* and *Faecalibacterium*, and is primarily associated with healthy individuals. Our results confirm that clustering based on the Bray–Curtis dissimilarity metric is primarily driven by species with high abundance and variance (Figure 4.C).

**Figure 4:**
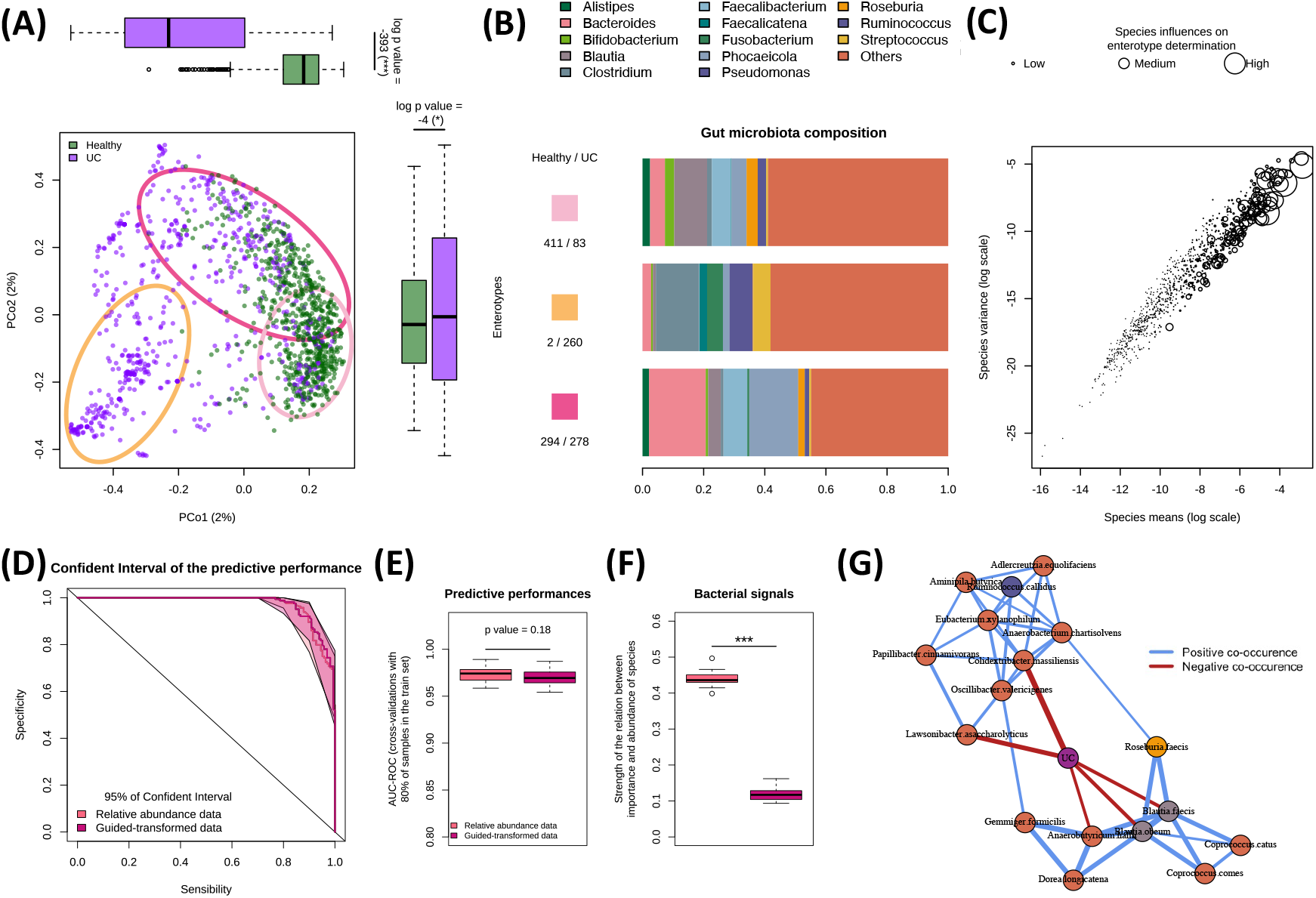
**(A)** Principal Coordinates Analysis (PCoA) with 80% confidence ellipses (CEs) for each cluster, represented by colored circles. **(B)** Bar plot of gut microbiota composition at the genus level, stratified by cluster. Only the 13 most abundant genera are shown. The yellow cluster is associated with the *Clostridium* enterotype, while the red cluster corresponds to the *Bacteroides* enterotype. **(C)** Abundance-variance relationship at the species level, illustrating the influence of each species on the clustering. Circle size reflects the species’ contribution to the clustering. **(D)** Predictive performance (with 95% confidence intervals) is compared between relative abundance and guided-transformed datasets. **(E)** Distribution of predictive performance across 20 cross-validation runs. **(F)** Distribution of *R*^2^ values across 20 cross-validations, indicating the strength of the relationship between species importance in the RF model and their abundances. **(G)** Co-occurrence network between the 20 most important bacterial species (as determined by the RF model trained on guided-transformed data) and host health status. Co-occurrences were evaluated using the Chi-square test (*χ*^2^); edge width is proportional to significance (lower p-values), with blue edges indicating positive associations and red edges indicating negative associations. The guided transformation has been applied to the relative abundance data.

We next compared the predictive performance of Random Forest (RF) models trained on relative abundance data and guided-transformed data derived from it. Each RF model was trained on 80% of the dataset, and predictions were generated on the remaining 20%. This process was repeated 20 times to assess performance stability by the distribution of AUC-ROC scores.

RF models trained on relative abundance data achieved a mean AUC-ROC of approximately 0.98. In particular, models trained on guided transformed data achieved the same performance (Figures 4.D-E), indicating that removing dominant bacterial signals did not affect predictive power. However, a comparison of the importance profiles of the features revealed that guided transformation alters the interpretability of the model. Indeed, when trained on guided transformed data, RF models are more dependent on minor bacterial signals (Figure 4.F). This observation is supported by lower values of *R*^2^ between species abundance and feature importance, indicating a shift away from high-abundance species (Figure 4.F).

The approach implemented here is closely related to out-of-sample deconfounding, providing empirical evidence that enterotype-related information may not be essential for predicting host health and can act as a confounding factor (*30*). The guided transformation instead allows the model to focus on minor bacterial signals that may carry greater relevance to the phenotype of interest.

Interestingly, while Wu et al. reported that *Ruminococcus gnavus* is more abundant in subjects with UC associated with ET-B and ET-C, our analysis suggests the presence of two distinct microbial networks that are not directly related to the pathology of UC, but can play protective roles (Figure 4G). The first group, located upstream of the UC node, includes *Ruminococcus callidus*, a species more abundant in healthy individuals. Although phylogenetically close to *R. gnavus. R. Callidus* exhibits a negative correlation in abundance with *R. gnavus*, suggesting competitive exclusion and a possible protective effect. Notably, *R. Callidus* has been suggested as a key bacterium that may protect against inflammatory bowel disease (*31*). In addition, as a fiber-degrading species, it belongs to the same group of beneficial microbes (including *Faecalibacterium prausnitzii*) that are reduced in children with pre-diabetes (*32*).

The second group, located downstream of the UC node, includes species consistently selected by RF models trained on relative abundance data. All samples expressing this group of species belong to healthy individuals, with no UC cases observed, further supporting a potential protective role (data not shown).

In summary, our results reinforce the notion that dominant bacterial signals can obscure minor but biologically relevant species. The application of guided data transformation not only preserves predictive performance but also enhances interpretability, offering novel insights into the complex interplay between microbial composition and host health.

## 3 METHODS

### 3.1 Study Populations

The present study used three distinct datasets of the gut microbiome to ensure a comprehensive analysis. The first dataset served as the reference dataset that we use for the demonstration and simulation sections (from 2.1 to 2.8). The remaining two data sets were used in the application sections (2.9 and 2.10) to illustrate the impact, effectiveness, and robustness of the proposed method.

#### 3.1.1 Inflammatory Bowel Disease (IBD) [reference dataset]

The reference data set used to observe the problems is registered in Bio-project n°PRJEB1220. (*18*) It consists of 396 samples, each characterized by the expression levels of 606 microbial species and an associated binary host health variable, 0 corresponding to a healthy sample (HLT) and 1 corresponding to a sample presenting an irritable bowel disease (IBD).

#### 3.1.2 Fat oxidation (FO) and insulin sensitivity [dataset n°2]

The second dataset is a small cohort (n = 50 and p = 248) where the binary variable of host health is the maximal oxidation of fat (FO) (*25*). It refers to the ability to oxidize fat during submaximal exercise in a fasted state, measured in g/min. The 0.4 g/min threshold for FO was selected because it has previously been identified as an effective cut-off to discriminate between populations with poor versus normal metabolic flexibility (*26, 27*). In this case, 0 corresponds to a healthy sample presenting an FO higher than 0.4, and 1 corresponds to a sample presenting an FO lower than 0.4.

#### 3.1.3 Ulcerative colitis (UC) [dataset n°3]

The third dataset comprises 12 publicly available datasets. All gut microbiomes have been curated in a previous study, provided in their Github: https://github.com/WXG920713/Gut-microbes. (*29*) They obtained gut microbiome metagenomic data from three sources: GMrepo, the European Bioinformatics Institute (EMBL-EBI), and Google Scholar, using specific search criteria related to ulcerative colitis (UC) and 16S rRNA gene sequencing. They generate a data set including 1328 samples that express 925 species. In this dataset, each sample is labeled with a binary host health variable, where 0 indicates a healthy individual (HLT) and 1 corresponds to a case of ulcerative colitis (UC).

### 3.2 Data transformation

#### 3.2.1 Relative abundance data

The non-transformed gut microbiome data corresponds to the compositional data. Compositional data are restricted within a simplex (*S*^*W*^), where data contain *D* parts of nonnegative numbers whose sum is 1. In this case, for each observation, the variables are limited to the interval [0, 1] (*33*).

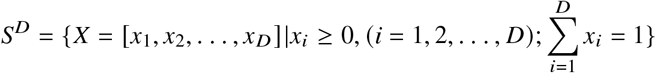

#### 3.2.2 Absence/presence data

The gut microbiome data is transformed into the absence/presence matrix, where 0 refers to the absence of a species in the sample, while 1 refers to the presence of the bacteria.

#### 3.2.3 Log transformed data

The log-ratio transformations eliminate the non-negativity constraint of compositional data and establish a one-to-one mapping onto real space, allowing researchers to use standard multivariate methods (*34*). In this paper, we focus specifically on the *centered* log*-ratio* (CLR) transformation due to its relevance in subsequent analyses. The CLR transformation uses the logarithm of the ratio of each component over the geometric mean of all components (*33*).

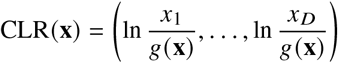

where *g*(*x*) is the geometric mean of the vector *x*, defined as:

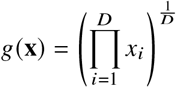

This transformation preserves the relative structure of the data while enabling robust statistical analysis.

### 3.3 Clustering

The clustering involves the following steps:

- The pairwise (dis)similarity or distance matrix is computed, denoted *D*.
- Hierarchical Agglomerative Clustering (HAC) is applied on *D*.
- The clustering is then obtained by applying the `cutree`function in R on the dendrogram derived from the HAC.
- The optimal number of classes is determined by selecting the number that maximizes the silhouette coeffcient.

#### 3.3.1 Assessment of clustering performance

To assess clustering performance, a confusion matrix is constructed between the host health status and the clusters. A *χ* ^2^ test is then applied to this matrix to evaluate the degree of association. The null hypothesis (*H*_0_) states independence between clustering and host health. The resulting p-values are log-transformed for interpretability. More negative log-transformed p-values indicate stronger statistical dependence between clusters and host health, thus reflecting improved clustering performance.

#### 3.3.2 Assessment of the species driving the clustering

To assess the contribution of minor versus dominant bacterial signals to the clustering, a Random Forest (RF) model is trained to predict cluster assignments. The mean abundances of the top 20 most important species, as determined by the RF model, are log-transformed. Lower values indicate that the clustering is predominantly driven by low-abundance bacterial species.

Additionally, the relationship between species abundance and their importance in the RF model is quantified by calculating the coeffcient of determination (*R*^2^). A high *R*^2^ value suggests that species with high abundance and variance exert a dominant influence on the clustering outcome, whereas a low *R*^2^ indicates a greater role for minor bacterial signals.

### 3.4 Predictive classification

#### 3.4.1 Assessment of predictive performance

The datasets under study are randomly partitioned into training and test sets. Machine learning models are fitted using the training data, and their predictive performance is evaluated on the corresponding test sets. Performance is assessed using the Area Under the Receiver Operating Characteristic Curve (AUC-ROC), averaged over 20 cross-validation iterations. Higher AUC-ROC values indicate superior classification performance in distinguishing host health status.

#### 3.4.2 Assessment of the species driving the classification

To evaluate the contribution of minor bacterial signals to the prediction, the mean abundances of the 20 most important species (identified via feature importance from the Random Forest (RF) model) are logtransformed. Lower log-transformed mean values indicate that the model’s predictions are primarily based on low-abundance (i.e., minor) bacterial species.

In addition, the relationship between species abundance and their predictive importance is quantified by computing the coeffcient of determination (*R*^2^). A high *R*^2^ value suggests that species with high abundance and variance predominantly influence the model’s predictions, whereas a lower *R*^2^ indicates a greater contribution from minor signals.

### 3.5 Numerical experiments and simulations

The dataset of the Inflammatory Bowel Disease is used to simulate the experimental data. The simulated host health (*Y*_sim_) is computed from both dominant and minor bacterial signals. We propose three scenarios in which the dependence of dominant bacterial signals on *Y*_sim_ is categorized as low, medium, or high. Those situations are tested in different dimensions where the number of samples varies (*n*_*sim*_ ∈ {75, 150, 300}). Fifty datasets have been simulated.

### 3.5.1 Simulation of the microbiome data

We propose a new method to simulate microbiome data with statistical characteristics similar to those of a reference dataset (especially the (dis)similarity between species, which can be related to their interaction).

Our method, described below, is inspired by the MIDAsim algorithm (*35*). MIDASim is a non-parametric approach that generally outperforms model-based simulation algorithms. However, in MIDASim, the size of the simulated sample is constrained to be the same as the size of the reference dataset. In our approach, we relax this constraint. Our novel algorithm has been implemented in the function `Simulator`in the Python module named `BiomeSampler`available at: https://github.com/pierrehouedry/BiomeSampler.

Let us denote by *X* = (*X*_*ij*_) ∈ ℝ^*n*^ ^×*p*^ our given data set of microbiome data and *X*_*sim*_ the simulated dataset. To simulate new data, we start by generating a binary sample that represents the absence-presence of the species in our dataset. This is done by first defining the absence-presence matrix P(*X*) = (*a*_*ij*_) ∈ ℝ^*n*^ ^×^ ^*p*^, where each entry *a*_*ij*_ is given by:

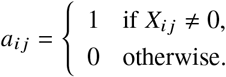

Once we have this binary matrix, we estimate its correlation matrix 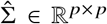. Given the numerical errors that may arise in the calculation, we need to ensure that ε is positive semi-definite to make it suitable for sampling from a multivariate normal distribution. To achieve this, we use Higham’s algorithm, which finds the closest positive semi-definite matrix (in the Frobenius norm sense) 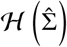.

Next, we simulate the absence-presence data *AP* by sampling from the multivariate normal distribution:

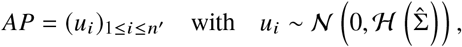

where *n* ^′^ is the desired number of samples.

To maintain a proportion of zeros similar to the original data, we define the simulated absence-presence matrix *Ā* = (*ā*_*ij*_) ∈ ℝ^*n*^ ′ ^× *p*^, where:

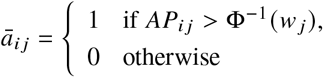

where Φ denotes the cumulative distribution function of the standard normal distribution 𝒩 (0, 1), and the estimated proportion of zeros for each species is calculated as follows:

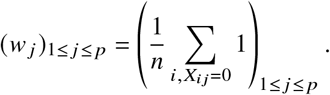

Once the absence-presence matrix is simulated, we move on to simulate the abundance values for the non-zero entries. The aim is to simulate according to the empirical law. For each feature 1 ≤ *j* ≤ *M*, let 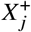 be the vector containing only the strictly positive values of (*X*_*ij*_)1 ≤ *i* ≤ *n*. To perform density estimation, we use a bandwidth, which is chosen adaptively for each feature using Silverman’s rule of thumb:

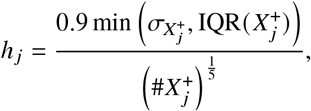

where 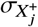 and IQR 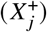 represent the standard deviation and interquartile range of 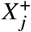, respectively.

We simulate the dataset 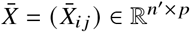 by setting:

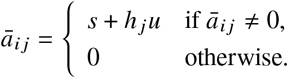

Here, *u* ~ 𝒰 ([0, 1]), and *u* is uniformly chosen from the elements of 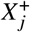. We summarize the method in the Supplementary materials in the Algorithm 2. The comparison of our method with other existing methods (*35, 36, 37*) is presented in Figure S3. Moreover, an example of a simulated dataset is presented in Figure 6.

### 3.5.2 Simulation of the host health

The simulated host health outcome, denoted*Y*_sim_, was modeled as a function of both dominant and minor bacterial signals. To investigate the influence of dominant bacterial signals, we constructed three scenarios in which these signals exerted low, medium, and high influence on *Y*_sim_. In these scenarios, the contribution of minor bacterial signals decreased proportionally. However, they remained relevant to the simulated phenotype.

Specifically, *Y*_sim_ was generated by combining two probabilities: (1) the probability that a sample exhibits a gut microbiota composition characterized by a high ratio of species from the *Prevotella* genus relative to the *Bacteroides* genus (*P*_1_), and (2) the probability that a sample presents with Irritable Bowel Disease (IBD) (*P*_2_). Each probability is estimated using a machine learning model trained on a reference dataset. Specifically, *P*_1_ is derived from a model fitted to highly abundant species to predict the first level of clustering (i.e., enterotype), while *P*_2_ is obtained from a separate model trained on low-abundance species to predict host health status. Both outputs are expressed as probabilities. The figure 5 and the algorithm 3 give details of the simulation of *Y*_sim_. In this context, the simulated host health was then defined as:

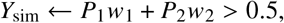

where *w*_1_ + *w*_2_ = 1 and *w*_1_ ∈ {0, 0.08, 0.15}. A higher value of *w*_1_ indicates a greater contribution of the dominant signal (*P*_1_) to *Y*_sim_. For example, when *w*_1_ = 0.15, IBD samples are more likely to be associated with clusters characterized by a high abundance of *Prevotella* species. We confirm these simulation plans in Figure 6.

**Figure 5:**
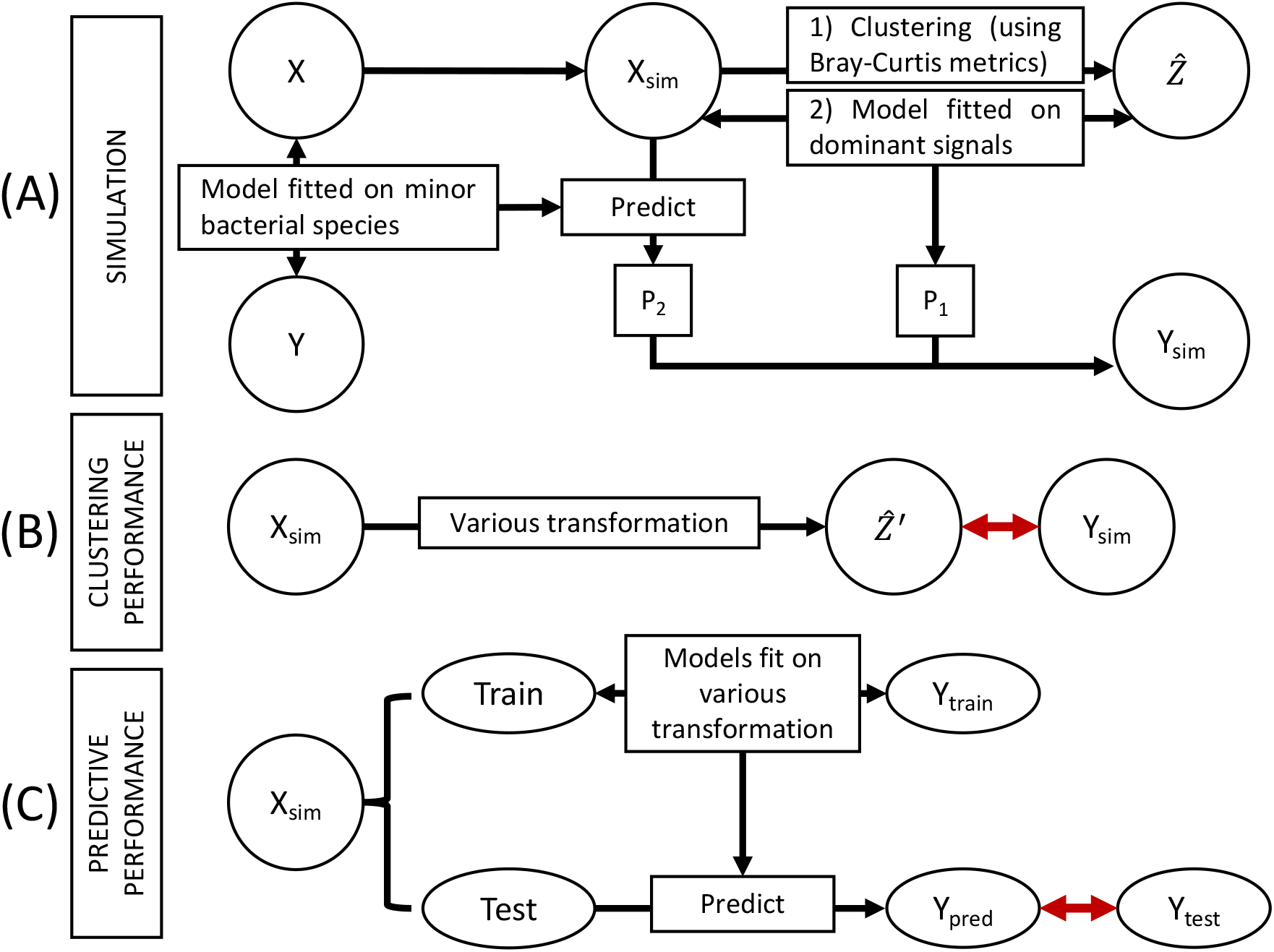
Design of numerical experiments. (**A**) The reference dataset (**X**) is used to simulate an experimental dataset (*X*_sim_). A predictive model is trained on minor bacterial species to estimate host health (*Y*) in the reference dataset. The probability of belonging to the IBD group (*P*_2_) is then computed based on this model. Clustering is performed on *X*_sim_ to infer enterotypes, denoted as 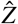. We consider three scenarios reflecting low, medium, and high dependence between 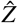 and *Y*_sim_, effectively introducing varying levels of noise into the host health signal. The noise is introduced as follows: (i) Fit a Random Forest on the dominant bacterial signals and then (ii) predict the probability of belonging to the *Prevotella* enterotype (denotes *P*_1_). (iii) Finally, a conditional probability is computed from *P*_1_ and *P*_2_ as it is described in the materials and methods. Noise was introduced through the following procedure: (i) a Random Forest classifier was trained on the dominant bacterial signals; (ii) this model was then used to estimate the probability of belonging to the Prevotella enterotype, denoted as *P*_1_; and (iii) a conditional probability was subsequently computed from *P*_1_ and *P*_2_, as detailed in the Materials and Methods section. These scenarios are evaluated across different sample sizes (*n*_sim_ ∈ {75, 150, 300}), with 50 datasets simulated for each condition. (**B**) Clustering performance is assessed under each experimental condition. Alternative clusterings 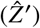 are obtained by applying HAC on various data transformations. A *χ*^2^ test is applied to the contingency table between 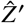 and *Y*_sim_ to quantify their association. This evaluation step is illustrated by a prominent red arrow in the figure. (**C**) Predictive performance is evaluated for each experimental condition. The dataset *X*_sim_ is split into training and testing subsets. Machine learning models are trained on the training data and evaluated on the test data. Predictive accuracy is assessed using the Area Under the Receiver Operating Characteristic Curve (AUC-ROC), based on the comparison between predicted and true host health outcomes (*Y* _pred_ vs. *Y*_test_). This evaluation step is also represented by a prominent red arrow in the figure.

**Figure 6:**
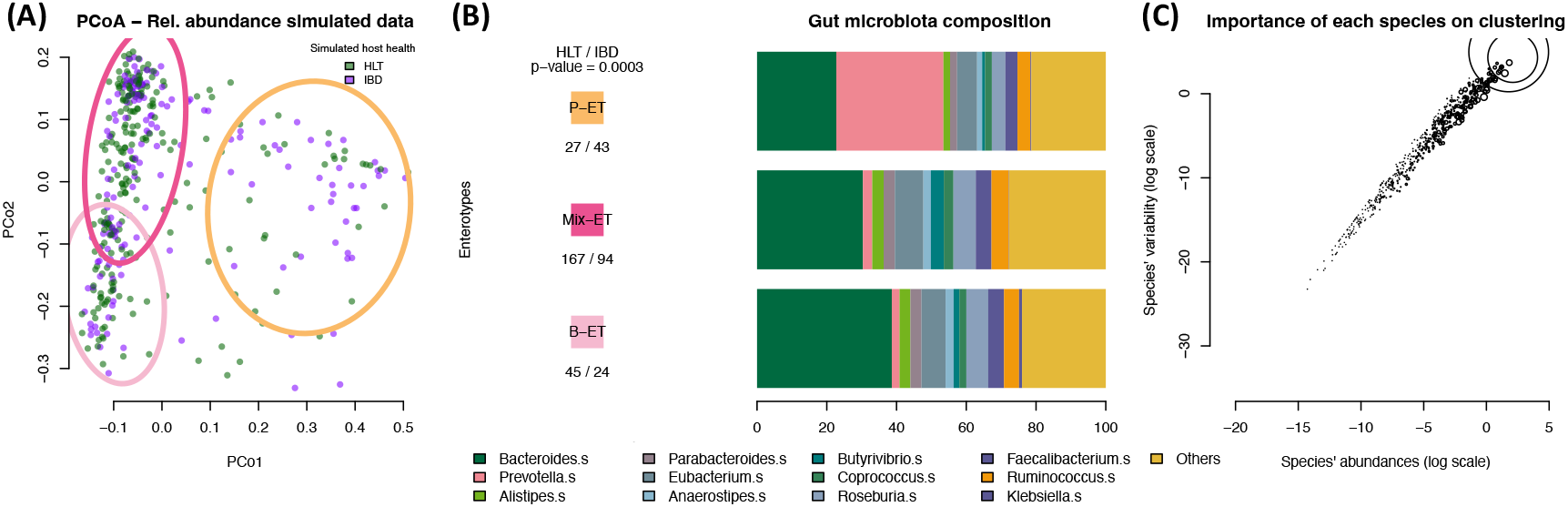
Example of data simulated from the reference dataset with *n*_*sim*_ = 300 **and** *w*_1_ = 0.15. **(A)** Principal Coordinates Analysis (PCoA) with 80% confidence ellipses (CEs) for each cluster, represented by colored circles. **(B)** Bar plot of gut microbiota composition at the genus level, stratified by cluster. Only the 13 most abundant genera are shown. **(C)** Abundance-variance relationship at the species level, illustrating the influence of each species on the clustering. Circle size reflects the species’ contribution to the clustering. We confirm that the simulation produced a configuration characterized by three distinct enterotypes, each driven by species predominantly from either the *Prevotella* or *Bacteroides* genus. As anticipated based on our simulation design, samples assigned to the enterotype dominated by *Prevotella* species were those most frequently associated with Inflammatory Bowel Disease (IBD). This association yielded a statistically significant result (p-value *<* 0.001), rejecting the null hypothesis (*H*_0_) that enterotype composition is independent of the simulated host health status

## Acknowledgments

The authors thank the anonymous reviewers for their valuable suggestions.

## Funding

This work is supported in part by funds from the University of Rennes and the French National Research Agency within the framework of the PIA France 2030 program for EUR DIGISPORT (ANR-18-EURE-0022) projects.

## Author contribution

DM and VM developed the conceptual framework, DM conceived the experiment(s), DM and PH conducted the experiment(s), PH conceived the simulation microbiome algorithm, DM, PH, and VM analyzed the results. DM, PH, FD and VM wrote and reviewed the manuscript.

## Competing interests

There are no competing interests to declare.

## Data and materials availability

The R scripts and the Python packages used for the simulation and the numerical experiments are available at: https://github.com/pierrehouedry/BiomeSampler. Any additional information required to reanalyze the data reported in this paper is available from the lead contact upon request.

## Notes

### Competing Interest Statement

The authors have declared no competing interest.

