## Supplemental Figures 1-3 for "A conceptual framework for revealing minor bacterial signals in microbiome data through guided data transformation"

**This PDF file includes:**

Materials and Methods
Figures S1 to S3

### Materials and Methods

#### 3.5.3 Algorithm of the microbiome simulation

The simulation of the microbiome algorithm is described here.

---

**Algorithm 2** Simulation of $X_{sim}$ from $X$

---

**Require:** $X$ the compositional data, $n'$ the desired number of samples

$\Sigma \leftarrow \text{corr}(\mathbb{1}_{x>0}(X))$.

$w \leftarrow \left(\frac{\#(X_j=0)}{n}\right)_{1 \le j \le p}$

Apply the Higham's algorithm to $\Sigma$ to get the closest semi-definite positive matrix $\mathcal{H}(X)$

$AP \sim \backslash(0, \mathcal{H}(\Sigma))$

$\bar{A} \leftarrow (AP > \Phi^{-1}(w))$

$X_j^+ \leftarrow X_j > 0$

**for** $j = 1, \ldots, p$ **do**

    $h_j \leftarrow \frac{0.9 \min\left(\sigma_{X_j^+}, \text{IQR}(X_j^+)\right)}{\left(\#X_j^+\right)^{\frac{1}{5}}}$

**end for**

**for** $i = 1, \ldots, n'$ **do**

    **for** $j = 1, \ldots, p$ **do**

        **if** $\bar{a}_{ij} \neq 0$ **then**

            $x \sim \mathcal{U}(X_j^+)$

            $u \sim \mathcal{U}([0, 1])$

            $\bar{x}_{ij} \leftarrow x + h_j u$

        **else**

            $\bar{x}_{ij} \leftarrow 0$

        **end if**

    **end for**

**end for**

**return** $(\bar{x}_{ij}) \in \mathbb{R}^{n' \times p}$.

---

#### 3.5.4 Algorithm of the host health simulation

The simulation of the host health algorithm is described here.

---

**Algorithm 3** Simulation of $Y_{\text{sim}}$ from $\mathbf{X}^{(m)}$

---

**Require:** $X$ the compositional data

$D = (d_{ij}) \in \mathbb{R}_+^{n \times n}$ (Bray-Curtis dissimilarity)

Apply HAC on D to estimate $Z^{(1)}$.

Select the species with the highest degree of information in $X$ and denote this community as dominant bacterial signals and fit a random forest model $\text{RF}_1$ on $\text{CLR}(X)$ to predict the enterotype labels.

Select the species with the lowest degree of information in $X$ and denote this community as minor bacterial signals and fit a random forest model $\text{RF}_2$ on $\text{CLR}(X)$ to predict the host health labels.

Simulate $X_{\text{sim}}$ with Algorithm 2

Apply Algorithm **??** on $X_{\text{sim}}$ to compute $r_{sim}$.

Apply $\text{RF}_1$ on $\text{CLR}(X_{\text{sim}})$ to estimate the probability $P_1$ of belonging to the *Prevotella* enterotype.

Apply $\text{RF}_2$ on $\text{CLR}(X_{\text{sim}})$ to estimate the probability $P_2$ of belonging to the IBD group.

Dependency between $Y_{\text{sim}}$ and both dominant bacterial signals and minor signal is described as follows:

```
for w1 ∈ {0, 0.08, 0.15} do
    if P1 w1 + P2 w2 > 0.5 then
        Y_sim ← IBD
    else
        Y_sim ← HLT
    end if
end for
return Y_sim
```

---

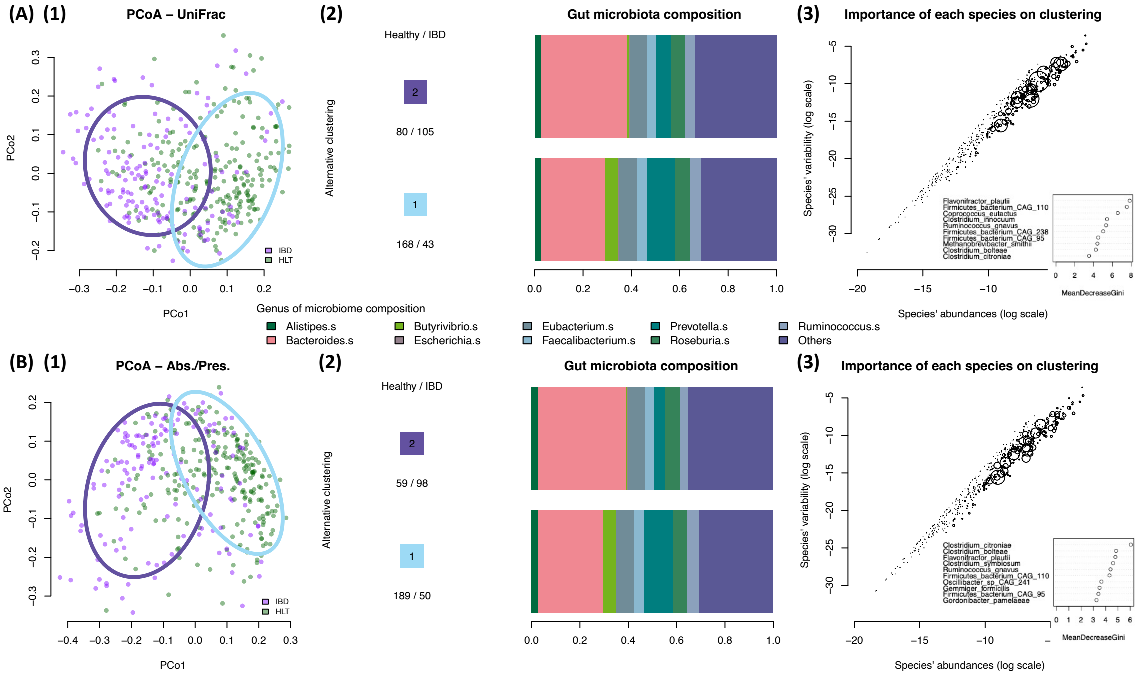


**Figure S1**: **Transformation of data improves the clustering performance.** (**1**) Principal Coordinates Analysis and the 80% confidence ellipse of the clusters (in colored circles) are presented. (**2**) Barplot of the microbiome composition, at the genus level, for each cluster. (**3**) The abundance/variance relationship at the species level. The influence of each species on the clustering (see Materials and Methods) is visually represented by the size of the circles in the abundance-variance relationship plot. The 10 most important species driving the clustering (i.e., the 10 largest circles) are shown with their corresponding mean decrease in Gini index. The following demonstration has been applied to both Absence/presence data (**A**) and Unifrac-based distance matrix (**B**).

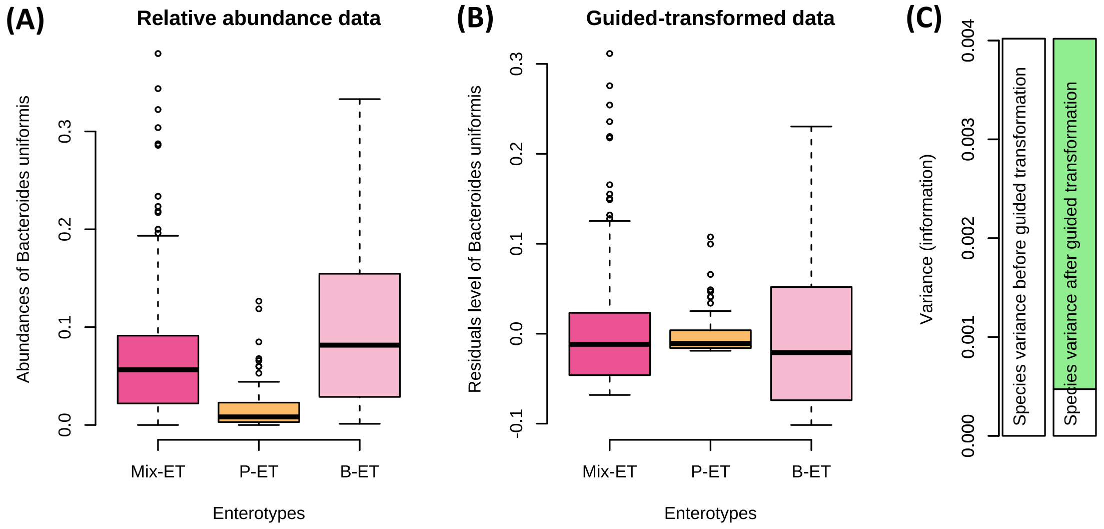


**Figure S2**: **Example of removing the information of the clustering related to the dominant bacterial signals in the dataset n°1.** (**A**) The abundance distribution of *Bacteroides Uniformis* depending of the first set of clusters ($\hat{Z}$). (**B**) The residual levels of *Bacteroides Uniformis* after the guided transformation. (**C**) The variance of *Bacteroides Uniformis* before and after the guided transformation. The part in light green refers to the information that is removed in the guided-transformed data. The part in white refers to the information in non-transformed data and transformed data, respectively, from left to right.

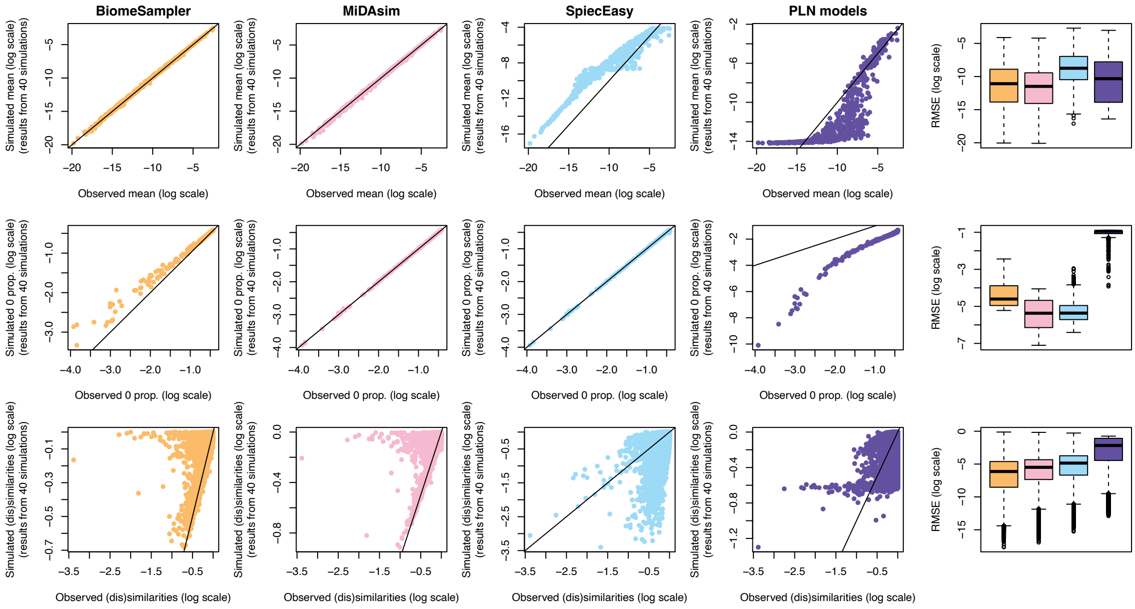


**Figure S3**: **Evaluation of the quality of simulation of four different methodologies.** The first four columns illustrate the linear relationship of specific parameters (mean, zero proportion, and interaction index) between the simulated and reference datasets, where each point represents a species. The greater the deviation of a point from the black reference line, the poorer the performance of the simulation for that species. The first row of plots displays the mean abundance per species, the second row shows the simulation performance concerning zero inflation (i.e., presence of zeros), and the third row presents the dissimilarity index across species. The final column shows the distribution of the root mean square error (RMSE) for each studied parameter (mean abundance, proportion of zeros, and interspecies interactions), where lower RMSE values indicate better simulation performance. The MiDAsim outperforms the algorithm that we developed (`BiomeSampler`) to simulate the zeros ($p < 0.001$), while the `BiomeSampler` has greater results in simulating the interaction between species ($p < 0.001$).
